# Tuning of ventral tenia tecta neurons of the olfactory cortex to distinct scenes of feeding behavior

**DOI:** 10.1101/455089

**Authors:** Kazuki Shiotani, Hiroyuki Manabe, Yuta Tanisumi, Koshi Murata, Junya Hirokawa, Yoshio Sakurai, Kensaku Mori

**Affiliations:** Laboratory of Neural Information, Graduate School of Brain Science, Doshisha University, Kyoto, Japan; Research Fellow of the Japan Society for the Promotion of Science, Tokyo, Japan; Division of Brain Structure and Function, Faculty of Medical Sciences, University of Fukui, Fukui, Japan; The University of Tokyo, Tokyo, Japan

## Abstract

Ventral tenia tecta (vTT) is a part of the olfactory cortex that receives both olfactory sensory signals from the olfactory bulb and top-down signals from the prefrontal cortex. To address the question whether and how the neuronal activity of the vTT is modulated by prefrontal cognitive processes such as attention, expectation and working memory that occurs during goal-directed behaviors, we recorded individual neuronal responses in the vTT of freely moving awake mice that performed learned odor-guided feeding and drinking behaviors. We found that the firing pattern of individual vTT cells had repeatable behavioral correlates such that the environmental and behavioral scene the mouse encountered during the learned behavior was the major determinant of when individual vTT neurons fired maximally. Furthermore, spiking activity of these scene cells was modulated not only by the present scene but also by the future scene that the mouse predicted. We show that vTT receives afferent input from the olfactory bulb and top-down inputs from the medial prefrontal cortex and piriform cortex.

These results indicate that different groups of vTT cells are activated at different scenes and suggest that processing of olfactory sensory information is handled by different scene cells during distinct scenes of learned feeding and drinking behaviors. In other words, during the feeding and drinking behavior, vTT changes its working mode moment by moment in accord with the scene change by selectively biasing specific scene cells. The scene effect on olfactory sensory processing in the vTT has implications for the neuronal circuit mechanisms of top-down attention and scene-dependent encoding and recall of olfactory memory.

## Introduction

In mammals, olfactory sensory information detected by sensory neurons in the olfactory epithelium is transmitted via the olfactory bulb to the olfactory cortex. Mitral and tufted cells in the olfactory bulb project axons directly to various areas of the olfactory cortex that includes anterior olfactory nucleus, ventral and dorsal tenia tecta, dorsal peduncular cortex, anterior piriform cortex, olfactory tubercle, posterior piriform cortex, nucleus of the lateral olfactory tract, anterior cortical amygdaloid nucleus, posterolateral cortical amygdaloid nucleus, and lateral entorhinal area (Neville and Haberly, 2004; Igarashi et al., 2012).

Despite the accumulation of knowledge about how odors are coded by olfactory sensory neurons (Buck and Axel, 1991) and how olfactory sensory signals are processed by neural circuits in the olfactory bulb and olfactory cortex (Mori and Sakano, 2011, Wilson and Sullivan, 2011, Mori et al., 2013), little is known about how olfactory cortical areas translate olfactory sensory information into behavioral responses (Choi et al., 2011). In this study, we focused on the ventral tenia tecta (vTT), an unexplored area of the olfactory cortex located at the ventromedial part of olfactory peduncle, and asked the question how the vTT translates odor signals from foods and environment into behaviors that are related to obtaining and consuming food and water.

vTT has a three-layered cortical structure (Haberly and Price, 1978, Brunjes et al., 2011). Principal neurons in the vTT are pyramidal cells that receive olfactory bulb inputs onto apical tuft dendrites in layer Ia, and Ib-association fiber inputs from other areas of the olfactory cortex. In addition, proximal apical dendrites and basal dendrites (in layers II and III) of vTT pyramidal cells receive deep association fiber inputs from the piriform cortex and top-down inputs from the medial prefrontal cortex (Luskin and Price, 1983; Hoover and Vertes, 2011). In this study, we demonstrate the connectivity pattern of vTT using a retrograde tracer. vTT receives inputs from the olfactory bulb, anterior piriform cortex, posterior piriform cortex, and medial prefrontal cortex. vTT massively projects axon to the olfactory bulb, anterior olfactory nucleus, and anterior piriform cortex.

Physiological studies of visual, auditory and somatosensory cortices showed that neurons of the neocortical sensory areas receive not only sensory signals from the external world but also top-down signals generated internally by higher level cognitive processes, including attention, expectation, working memory and decision making (Gilbert and Sigman, 2007; Roelfsema and deLange, 2016). In the olfactory cortical areas, olfactory tubercle neurons represent goal-directed behaviors and show enhanced odor responses when rats selectively direct attention to odors (Gadziola & Wesson, 2016; Carlson et al., 2018) and c-fos activity of these neurons changes with different motivated behaviors (Murata et al., 2015).

We therefore supposed that neurons in the vTT not only receive olfactory sensory inputs from the olfactory bulb but might be also influenced by top-down inputs generated in association with higher cognitive processes that are necessary for performing goal-directed behaviors including feeding and drinking behaviors (Bushman and Miller, 2014).

To address the question whether vTT neurons receive top-down signals in association with higher level cognitive processing, we recorded spiking activity of vTT neurons during odor-guided feeding and drinking behaviors. We trained mice to perform two types of odor-guided behaviors in two tasks. One group of mice were trained to associate an odor (either eugenol or vanilla essence) with sugar reward. We also trained these mice to associate a different odor (almond essence) with aversive consequences after sugar eating, i.e., intraperitoneal injection of lithium chloride (Raineki et al., 2009). The other group of mice were trained to associate an odor (eugenol) in the odor port with the appearance of water reward in the reward port that is located at the left of the odor port. These mice were trained to associate a different odor (amyl-acetate) in the odor port with no-reward in the reward port.

Analysis of firing pattern of vTT neurons during the feeding and drinking behaviors showed clear tuning of individual neurons to distinct scenes (i.e., distinct environmental and behavioral contexts) of learned behaviors. The results indicate that the function of vTT is not fixed but changes moment by moment in a scene-dependent manner during the whole sequence of feeding and drinking behaviors.

## Results

### Scene specific activity of vTT cells during the odor-guided eating or no-eating task

Mice were trained to perform an odor-guided behavioral task that required decision making between eating or no-eating based on the presented odor cue (Fig. 1a). We randomly presented sugar on a dish with one of three different cue odors (eugenol, vanilla essence, or almond essence) at an arbitrary position in the test cage. In the training sessions, eugenol odor and vanilla odor were associated with sugar reward, whereas almond odor was associated with sugar and aversive consequence (LiCl injection). After the learning, mice showed high accuracy rate (> 0.8) of eating or no-eating behavioral response to the cue odor throughout the session (Fig. 1b, 5 trials/block, average for 63 sessions from 6 mice). We presented also powder chow on a dish in trials that were randomly inserted among the above trials.

**Figure 1:**
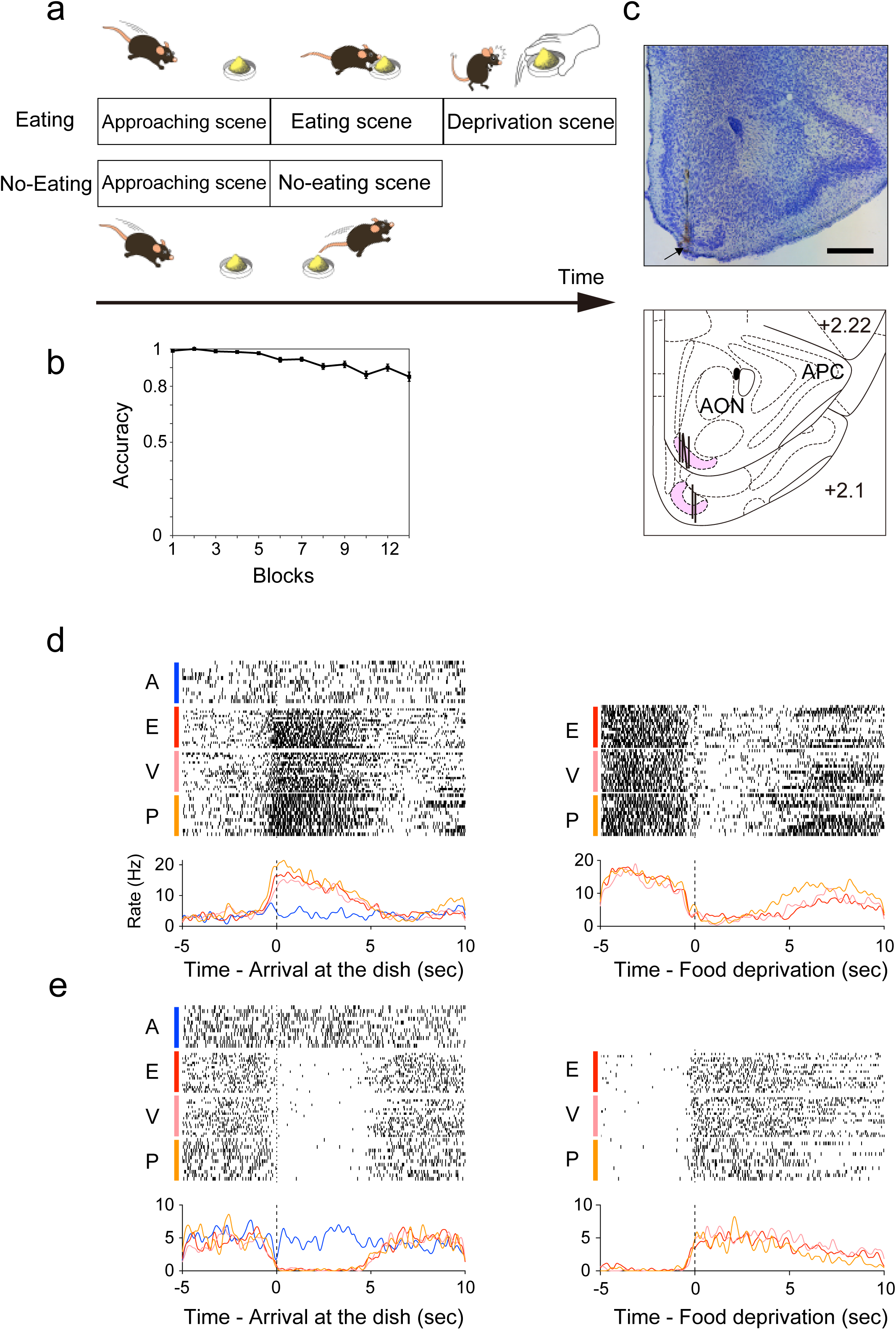
Firing pattern of vTT neurons during odor-guided eating or no-eating task and food deprivation. (a) Scene development during odor (eugenol, vanilla essence, powder chow)-guided eating task (upper illustration) and odor (almond oil)-guided no-eating task (lower illustration). Black arrow indicates time axis. Scenes develop with time from left to right. (b) Behavioral accuracy of the odor-guided eating or no-eating task (5 trials / block, n = 6 mice). (c) Top: Histological identification of recording sites. Nissl stained section of the olfactory peduncle. An arrow indicates electric lesion of recording site in the vTT. Scale bar, 500 μm. Bottom: Recording tracks in the vTT. Pink area shows vTT. Vertical thick lines indicate recording tracks. APC, anterior piriform cortex; AON, anterior olfactory nucleus. (d) Firing pattern of an eating scene cell during the odor-guided eating or no-eating task. Raster plots and peri-event time histogram (PETH) were aligned at the time when the mouse arrived at the dish (vertical dashed line at 0 in the left PETH) or at the time when the food was deprived (vertical dashed line at 0 in the right PETH). A, almond oil odor and no-eating (blue); E, eugenol odor and eating (red); V, vanilla essence odor and eating (pink); P, powder chow odor and eating (yellow). (e) Firing pattern of an instrumental scene cell. Raster plots aligned at the arrival (left PETH) and food deprivation (right PETH).

When the food dish was presented, the mouse approached the dish, and upon arrival at the dish the mouse showed either eating behavior or no-eating behavior depending on the odors attached to the food dish (Fig. 1a). We defined approaching scene as the time window between the start of approach behavior and the arrival at the food dish. We also defined eating scene as the time window between the arrival at the food dish and the end of eating, and no-eating scene as the time window between the arrival at the dish and 10 sec after the arrival during which the mouse did not eat any food. Average duration of the approaching scene was 3.5 sec in case of odor-guided eating behavior and 2.9 sec in case of odor-guided no-eating behavior. At the timing about 6.0 sec after the mouse started to eat in eating trials, we suddenly deprived the food dish even though the mouse was in the middle of eating. We defined deprivation scene as the time window between the moment of the food deprivation and 5 sec after the deprivation.

We measured spiking activity of individual vTT cells in six mice using extracellular tetrode recordings while the mice performed the eating and no-eating tasks (Fig. 1c). To examine the behavioral correlate of vTT cell firing pattern, we first selected vTT cells whose average firing rate during the trials was greater than 0.3 Hz for further analysis (n = 391 cells in total 63 sessions).

To assess firing pattern change of the vTT cells during the scene development of the eating task and no-eating task, we calculated peri-event time histograms (PETH) of the spiking activity of individual vTT cells with reference to the timing of arrival at the food dish or the timing of food deprivation. We compared the firing pattern of 391 vTT cells and noticed two types of cells that showed firing pattern change in opposite directions during the scene development.

One type of vTT cells were characterized by their increased spiking rate during the eating scene (Fig. 1d). These cells increased their spiking activity when the mouse arrived at the dish and started to eat sugar or powder chow and continued the increased spiking activity during the eating scene until the food deprivation (Fig. 1d left). These cells rapidly decreased the spiking activity when the food dish was deprived (Fig. 1d right). Because the maximal firing of these cells occurred during the eating scene and such high-frequency firing was absent in the absence of eating scene in no-eating task, we called these cells eating scene cells. A majority of eating scene cells fired maximally during the eating scene regardless of the odor type (eugenol odor, vanilla essence odor or powder chow odor) used for food cue and regardless of the taste of food (sugar or powder chow).

Another type of vTT cells showed decreased firing rate during the eating scene (Fig. 1e). These cells decreased spiking activity upon the arrival at the dish (Fig. 1e, left), and the suppression of spiking activity continued during the eating scene until the food was deprived (Fig. 1e, right). In response to the food deprivation, these cells rapidly increased the spiking activity. Because the maximal firing of these cells occurred during either the approaching scene or the food deprivation scene, we called these cells instrumental scene cells.

Although the food dish was presented at a randomly selected place in the cage in each trial, the eating scene cells fired maximally whenever and wherever the mouse ate the food, suggesting that the maximal firing of these vTT cells relate to the eating scene but not to the mouse’s place in the cage (c.f. hippocampal place cells, O’Keef, 2007).

To classify the firing patterns of vTT cells during the scene development, we used principal component analysis (PCA) of the firing pattern of 391 vTT cells followed by unsupervised, hierarchical clustering (Fig. 2a). This analysis showed two major clusters of cells that were separated according to the magnitude of firing in distinct scenes (Fig. 2b). Cells in type 1 cluster were eating scene cells showing maximal firing during eating scene (Fig. 2a light blue lines in the dendrogram and 2b top). Two hundred and twelve vTT cells (54.2 %) were classified as type 1 cluster eating scene cells using this clustering method. We observed that many of these eating scene cells began to increase their firing rate before the mouse touched the food dish, indicating that the firing rate increase of these eating scene cells during the pre-touch period was not caused by actual food intake (Fig. 2b top).

**Figure 2:**
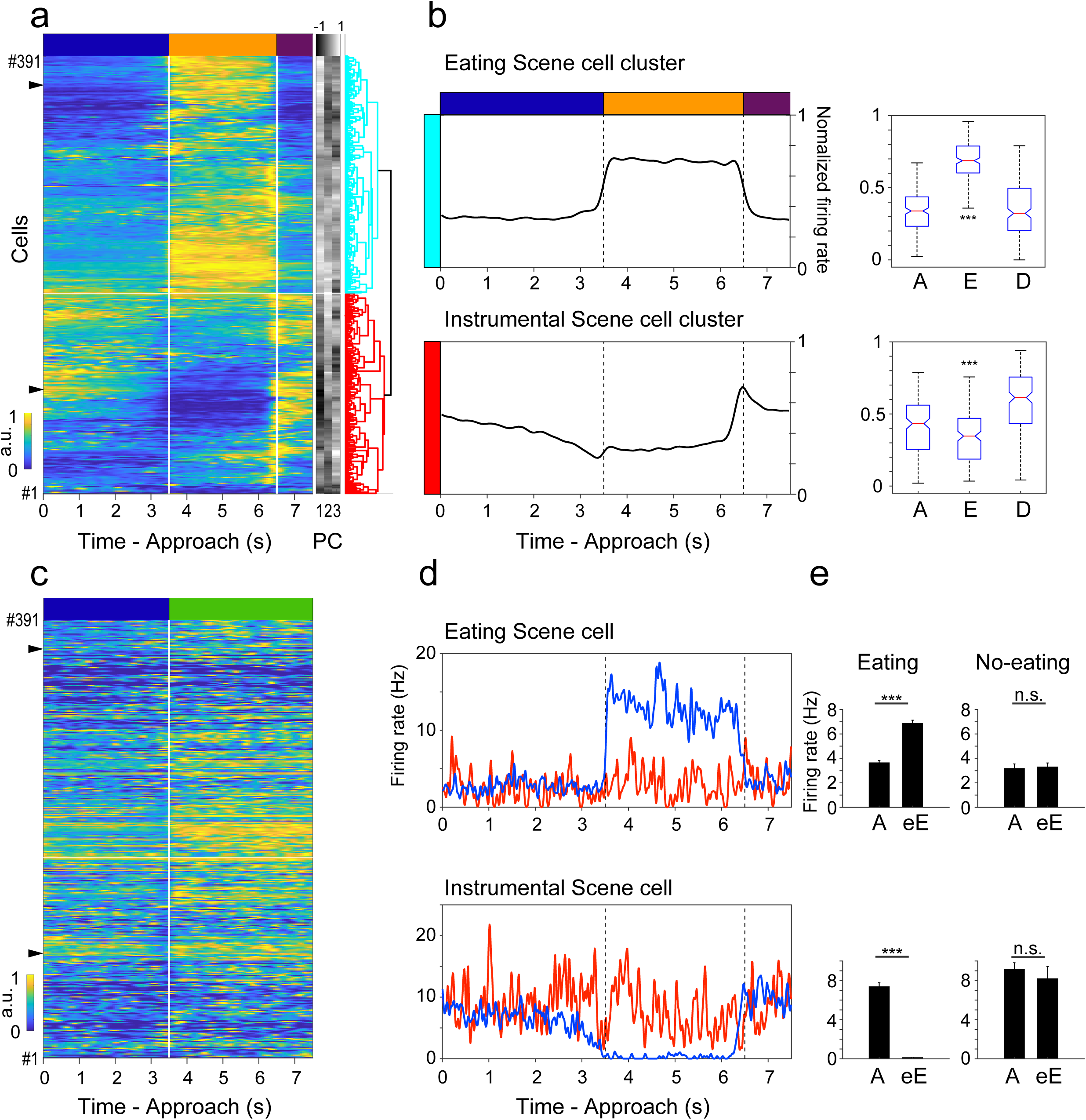
Pattern of firing rate change of vTT neurons along the time course of scene development. (a) Classification of a majority of vTT neurons into eating scene cells and instrumental scene cells based on principal component analysis (PCA) of their firing patterns. Left: Change in normalized firing rate along the time course of approaching scene (dark-blue bar), eating scene (orange bar) and deprivation scene (purple bar) during the go trials of odor-guided eating task. Each row represents one cell (cell #1 – cell #391). Time 0 indicates the start of approaching behavior toward the dish. The firing rate changes were aligned with the timing of start of approaching behavior (at 0 sec), arrival at the dish (at 3.5 sec) and the onset of food deprivation (at 6.5 sec). Duration of approaching scene from the onset of approaching behavior to the arrival at the dish) varied among different trials but is normalized in this graph to 0 - 3.5 sec. Duration of eating scene (from the start of eating to the onset of food deprivation) varied among different trials but is normalized to 3.5 - 6.5 sec. Arrows indicate representative cells showing in (d). a.u.: arbitrary unit of average normalized firing rate (0: minimal; 1: maximal) Right: Each row shows the first three principal components (1, 2, and 3) of the firing pattern of individual vTT cell. These values were used for the unsupervised hierarchical clustering, as shown in the right dendrogram. Two main clusters are shown by light-blue and red. (b) Left: Time course of the change in average normalized firing rate of cells in eating scene cell cluster (top) and that of cells in instrumental scene cell cluster (bottom) during the scene development. Right: Comparison of the average normalized firing rate of eating scene cell cluster (top) and instrumental scene cell cluster during the approach scene (left), eating scene (center) and deprivation scene (right). For all box plots, the central mark is the median, the top and bottom edges of the box are the 75th and 25th percentiles, and the whiskers are drown to the furthest observations. (Top, F(2, 633) = 368.57; bottom, F(2, 534) = 78.92, one-way ANOVA followed by post hoc Tukey test, ***, P < 0.01). A, approaching scene; E, eating scene; D, deprivation scene. (c) Change in the normalized firing rate along the time course of approaching scene (dark-blue bar) and no-eating scene (green bar) during the almond odor-guided no-eating task. Cells (#1 - #391) are the same cells that are shown in (a), and the order of neurons in the row is same as in (a). Arrows indicate representative cells showing in (d). (d) Firing pattern of a representative eating scene cell (top) and a representative instrumental scene cell (bottom) during the odor-guided eating task (blue line) and the no-eating task (red line). (e) Comparison of the average firing rate of late approaching scene (from 1.5 to 3.5 sec) and early eating scene (from 3.5 to 5.5 sec) during eating task (left) and no-eating task (right) in an eating scene cell (top) and an instrumental scene cell (bottom). A, late approaching scene; eE, early eating scene, ***, P < 0.01, unpaired t-test.

Cells in type 2 cluster were instrumental scene cells showing increased firing rate during the approaching and food deprivation scenes and decreased firing rate during the eating scene (Fig. 2a red lines, 2b bottom). These instrumental scene cells rapidly increased the firing rate around the timing of food deprivation. One hundred and seventy-nine vTT cells (45.8 %) were classified as type 2 cluster instrumental scene cells.

Many instrumental scene cells began to decrease their firing rate before the mice touched the food dish (Fig. 2b bottom), indicating that the firing rate decrease during the pre-touch period was not due to sensory inputs caused by actual food intake. Furthermore, these cells suddenly increased their firing rate before the food was deprived (Fig. 2b bottom), indicating that the firing rate increase during the pre-deprivation period was not caused by actual food deprivation.

These eating scene cells and instrumental scene cells showed only a minor change in firing rate when the mouse detected learned aversive odor (almond) and did not show eating behavior (Fig 1d, e, and Fig 2c, and d). To quantify the firing rate change of the scene cells at the transitions from approach scene to eating scene and from approach scene to no-eating scene, we aligned the firings of vTT cells in reference to the timing when the mouse arrived at the dish and examined the attached odor (3.5 sec on average after the start of approach behavior). We compared the average firing rate during a late approaching scene (from 1.5 to 3.5 sec on average after the start) with that during an early eating scene (from 3.5 to 5.5 sec on average after the start) in each scene cell (Fig. 2e). Forty-three percent of eating scene cells showed a significant increase of average firing during the early eating scene (E in Fig. 2e, upper left graph) compared with that during the late approaching scene (A in Fig. 2e, upper left graph) in eating trials. In the absence of eating scene in the no-eating trials, the average firing rate of these cells showed no significant change during an early no-eating scene (from 3.5 to 5.5 sec on average after the start of approach) compared with that during the late approaching scene (Fig. 2d top and 2e, upper right graph).

Sixty-nine percent of instrumental scene cells showed a significant decrease in average spiking activity in an early eating scene compared with the late approaching scene in eating trials, whereas the firing rate of these cells showed no significant change in the early no-eating scene in no-eating trials (Fig. 2d bottom and 2e, lower graphs). These results indicate that eating scene cells and instrumental scene cells receives nearly opposite influences at the transition from approach scene to eating scene, and that this influence is absent at the transition from approach scene to no-eating scene.

Although individual scene cells showed maximal firing rate in a specific scene, each scene cell showed a variety of firing pattern within the scene (Fig. 2a). To compare the firing profile of recorded scene cells, we aligned vTT cells by the timing of maximal firing as a function of scene development (Fig. 3). We found that individual vTT scene cells were tuned to a smaller scale scene or sub-scene within the eating scene or the instrumental scene. For example, some approaching scene cells were tuned to the early part of the approaching scene, while other approaching scene cells showed maximal firing rate at the late part of the approaching scene. A subset of eating scene cells showed maximal firing rate at the initial part of the eating scene, whereas another subset of eating scene cells were tuned to the middle part of the eating scene and the third subset of eating cells to the end part of the eating scene just before the food deprivation. We noted also the possibility that the recorded vTT cells represent all the scenes and sub-scenes that develop during the odor-guided eating task, which prompted us to define scenes and sub-scenes in more detail.

**Figure 3:**
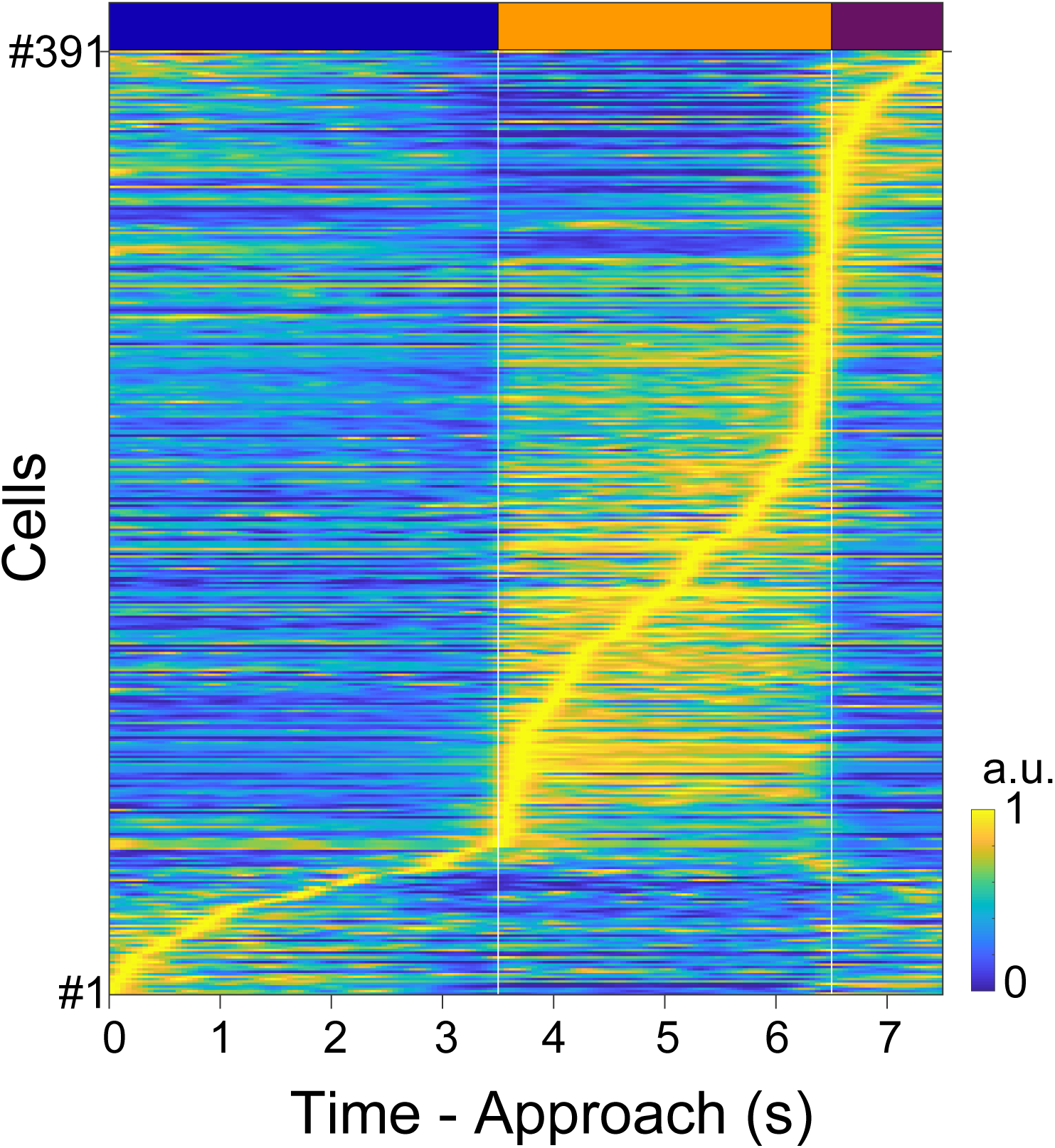
Alignment of vTT cells (#1 - #391) by the timing of maximal firing as a function of scene development during the odor-guided eating task. Top: dark-blue bar, approaching scene; orange bar, eating scene; purple bar, deprivation scene.

### Scene specific activity of vTT cells during the odor-guided Go/No-go task

To examine in more detail the scene- and sub-scene- dependency of maximal firing of individual vTT cells, we planned an odor-guided Go/No-go task to obtain water reward (Fig. 4a), in which we were able to define precisely the time window of approaching scene, odor checking scene, moving scene, waiting scene and water drinking scene. In this task, illumination of light at the right odor port instructed the mouse to start the task and approach to and nose poke into the odor port (approaching scene). Starting at the moment of the nose poke, one of cue odors was presented for 500 ms in the odor port. The mouse was required to sniff the cue odor and then keep nose poking for 500 ms after the cessation of odor stimulation. At 1 second after the onset of odor stimulation, the light was turned off and the mouse could withdraw its nose from the odor port. The period between the nose poke into and nose withdrawal from the odor port was defined as odor checking scene.

**Figure 4:**
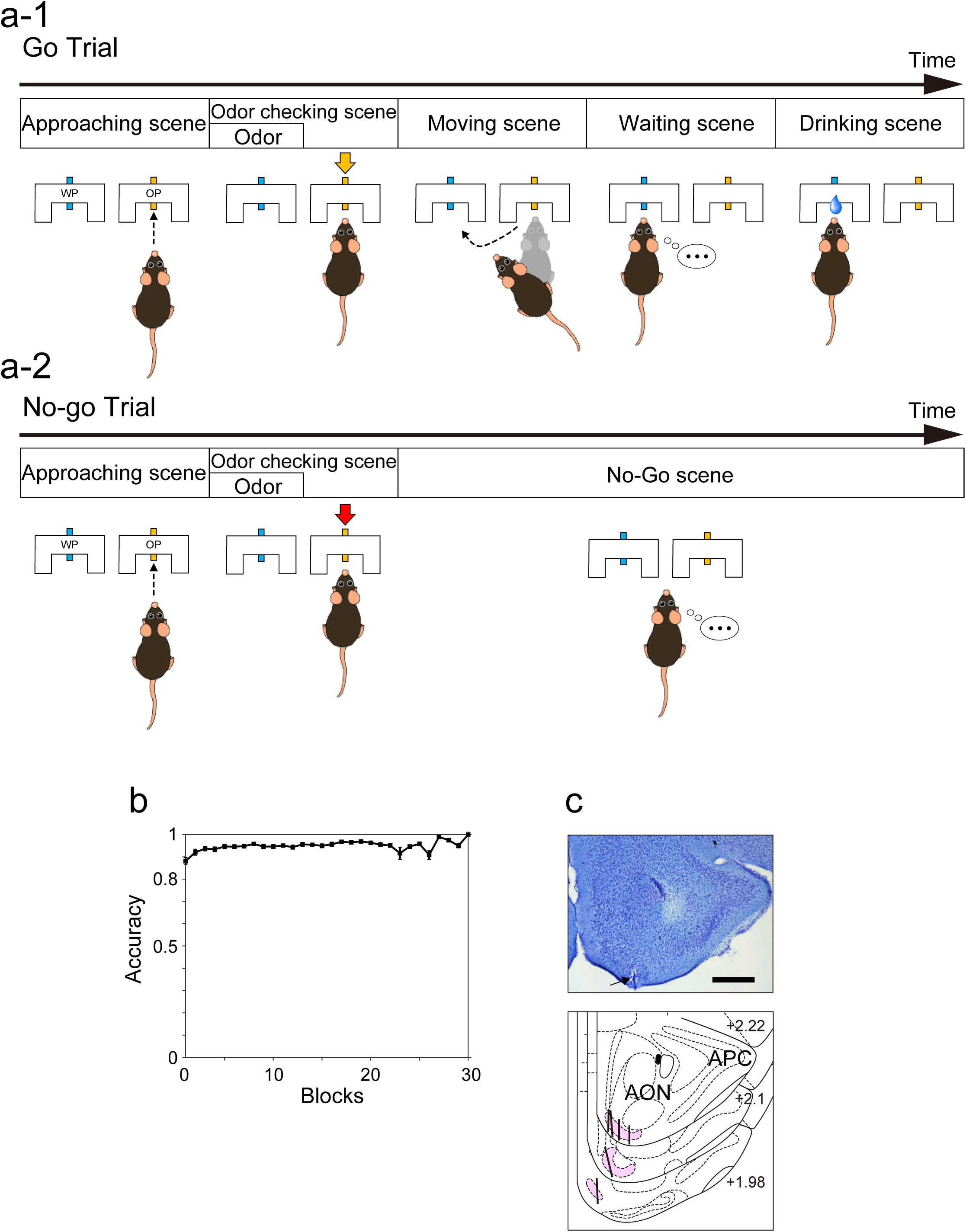
Odor-guided Go/No-go task to obtain water reward. (a-1) (Go Trial) Scene development during odor-guided go task. Eugenol odor was used as a cue for the go and drink task. Sequence of scenes invariably observed in this task was approaching scene, odor checking scene, moving scene, reward waiting scene, and drinking scene. Thick black arrow indicates time axis. Scenes develop with time from left to right. WP, water port; OP, odor port. Orange tube in the OP indicates an odor delivery tube. Orange arrow indicates eugenol odor delivery. Blue tube in the WP indicates a water delivery tube, and blue droplet in the drinking scene indicates water delivery. (a-2) (No-go Trial) Scene development during the task of odor guided no-go and wait trial. Amyl acetate odor was used as a cue for no-go and wait task. Red arrow indicates amyl acetate odor delivery. (b) Success rate of odor guided Go/No-go task (20 trials / block, n = 6 mice). (c) Histological identification of recorded sites. Arrow indicates electric lesion of recording site in the vTT. Scale bar, 500 µm. Thick lines indicate recording tracks in the vTT. Pink area shows vTT.

If go-cue odor (eugenol) was presented, the mouse was required to move to and poke its head into the left water port within 2 sec to obtain water reward. The period of moving from the odor port to the water port was defined as moving scene. At the water port, the mouse was required to keep its head in the port for 300 msec to wait for water delivery (waiting scene). Three hundred msec after the head poke, a drop of water (6 μl) was delivered (Fig 4a-1). Drinking scene was defined as the 1.2 sec period from the start of water delivery.

If no-go-cue odor (amyl acetate) was presented, the mouse was prohibited from poking its head into the water port for 2 sec after the end of odor delivery (No-go scene) (Fig. 4a-2). After mice were well trained, the behavioral accuracy kept more than 80% in a block (20 trials / block) throughout a session (average for 57 sessions from 6 mice, Fig. 4b).

We recorded the spiking activity of a total of 346 vTT cells from six mice using tetrodes while mice performed the odor-guided Go/No-Go task (Fig. 4c). To examine the relation of firing rate change of individual vTT cells to the development of behavioral scenes, we first selected a total of 288 vTT cells whose average firing rate during the trials was greater than 0.3 Hz for further analysis.

To classify the firing patterns of vTT cells during the scene development in the go trials, we used PCA of the firing pattern of vTT cells followed by unsupervised hierarchical clustering (Fig. 5a). This yielded six major clusters of cells. Four clusters of them showed highly increased firing rate in one or two specific scenes (Fig. 5a, b, light blue, red, dark green and pink) whereas two clusters showed highly increased firing rate in more than two scenes (Fig. 5a, b, yellow and light green).

**Figure 5:**
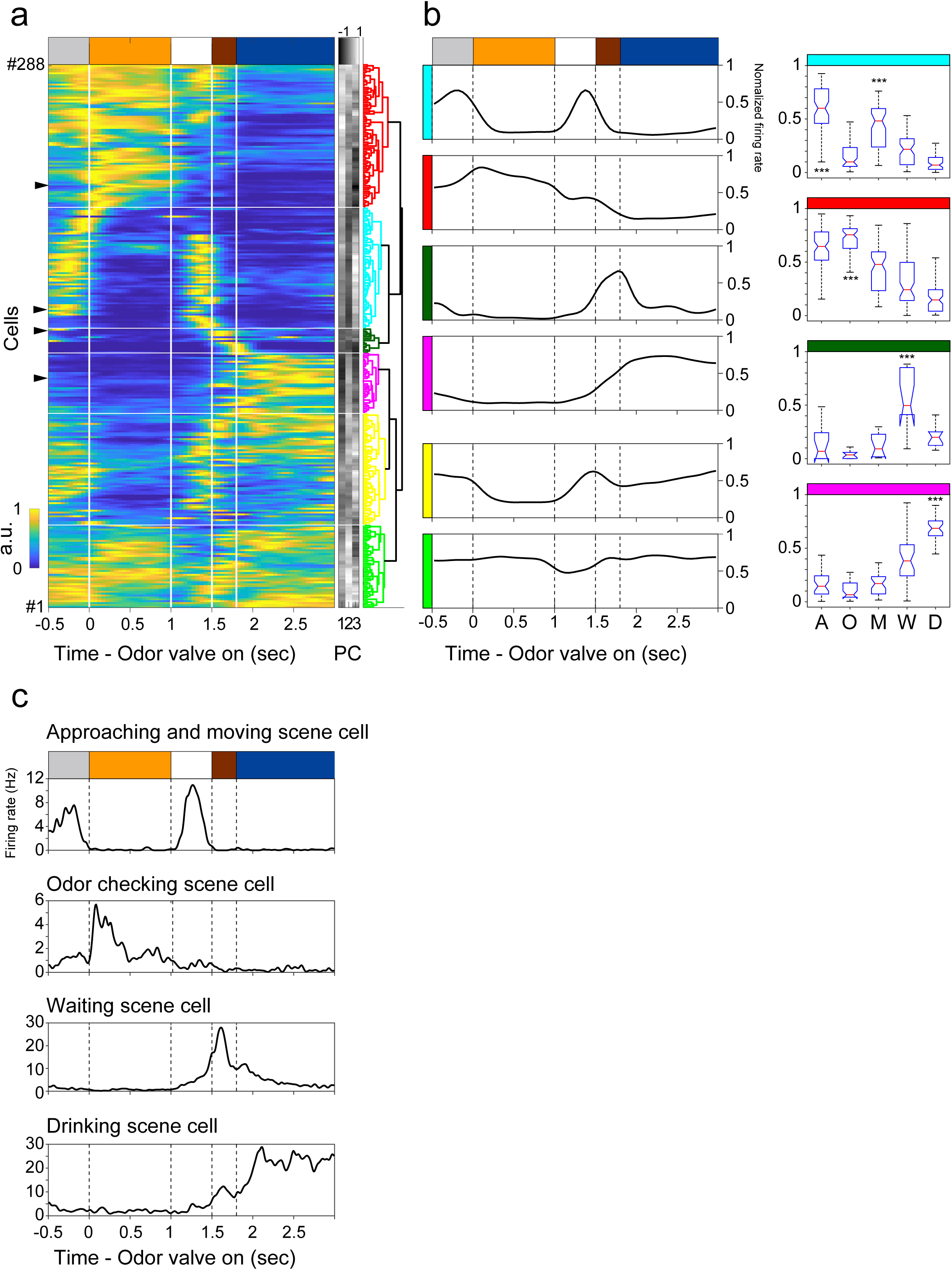
Firing pattern of vTT cells during odor-guided go trials. (a) PCA-based classification of vTT cells into four scene-specific clusters and two other clusters. Left: Firing rate change along the time course of approaching scene (gray bar), odor checking scene (orange bar), moving scene (white bar), waiting scene (brown bar), and drinking scene (dark-blue bar) during the odor-guided go trials. Each row represents one cell (cell #1 – cell #288). Time 0 indicates the timing of odor valve opening (start of odor delivery). The firing rate changes were aligned with the timings of onset of odor presentation (at 0 sec), withdrawal of nose from the odor port (at 1 sec), poking the mouth into the reward port (at 1.5 sec) and onset of water reward presentation (at 1.8 sec). Durations of approaching scene and moving scene varied among different trials but are normalized to −0.5 - 0 sec and 1 - 1.5 sec, respectively. Right: Each row represents the first three principal components (PC) (1, 2, and 3) of the firing pattern of an individual vTT cell. These values were used for the unsupervised hierarchical clustering, as shown in the right dendrogram. Four scene-specific cell clusters are shown in different colors (light blue, approaching and moving scene cell cluster; red, odor checking scene cell cluster; dark green, waiting scene cell cluster; pink, drinking scene cell cluster). Two unaccountable cell clusters are shown by yellow and light green. Arrows indicate representative cells showing in (c). (b) Average firing pattern of the vTT scene cells during the task. Color in left bar indicates the cell cluster shown by the same color in (a). Right: Comparison of the averaged normalized firing rate of approaching and moving scene cell cluster (light blue), odor checking scene cell cluster (red), waiting scene cell cluster (dark-green), and drinking scene cell cluster (pink) during each scene. Other two cell clusters with multi-scene activity are shown by yellow and light green. For all box plots, the central mark is the median, the top and bottom edges of the box are the 75th and 25th percentiles, and the whiskers are drown to the furthest observations (approaching and moving scene cell cluster, F(4,315)=103.23; odor checking scene cell cluster, F(4,375)=120.92; waiting scene cell cluster, F(4,60)=20.66; drinking scene cell cluster, F(4,155)=91.85, one-way ANOVA followed by post hoc Tukey test; ***, P < 0.01). A, approaching scene; O, odor checking scene; M, moving scene; W, waiting scene; D, drinking scene. (c) Firing pattern of a representative approaching- and moving-scene cell, an odor checking scene cell, a waiting scene cell, and a drinking scene cell along the time course of the go trials.

In the PCA of firing pattern of 288 vTT cells, 64 cells (22.2 %) were sorted into approaching and moving scene cell cluster (light blue cluster in Fig. 5a, b). Average firing rate of cells in this cluster was significantly higher in both the approaching scene and moving scene. A typical example of cells in this cluster is shown at the top histogram of Fig. 5c. This cell showed maximal firing during moving scene and high firing rate during approaching scene, whereas it showed diminished spiking activity during odor checking scene, waiting scene, and drinking scene.

The PCA sorted 76 cells (26.4 %) into odor checking scene cell cluster, and average firing rate of cells in this cluster was maximal in the odor checking scene (red cluster in Fig. 5a, b, an example is shown at the 2^nd^ histogram from the top in c). The firing rate of the odor checking scene cells was lower during the drinking scene.

The PCA sorted 13 cells (4.5 %) into waiting scene cell cluster, and average firing rate of cells in this cluster was maximal in the waiting scene (dark green cluster in Fig. 5a, b, an example is shown at the 3^rd^ histogram from the top in Fig. 5c).

The PCA sorted 32 cells (11.1 %) into drinking scene cell cluster. The average firing rate of cells in this cluster was maximal in the drinking scene (pink cluster in Fig. 5a, b, an example is shown at the bottom histogram in Fig. 5c). These drinking scene cells showed diminished spiking activity during odor checking scene. Thus odor checking scene cells and drinking scene cells showed firing rate change in nearly opposite directions during the scene development.

As shown in Fig. 5a-c, a majority of odor checking scene cells (red) began to increase the firing rate before the start of the odor checking scene, i.e., before the mouse poked its nose into the odor port and smelled the cue odor. This observation indicates that the firing rate increase in odor checking scene cells before the start of odor checking scene is not caused by the olfactory sensory input of the delivered cue odor.

Many waiting scene cells (dark green) began to increase the firing rate before the start of the waiting scene, i.e., before the mouse poked its mouth into the reward port, suggesting that the firing rate increase in waiting scene cells just before the waiting scene is not due to odors from the inside of the reward port. Furthermore, many drinking scene cells (pink) began to increase their firing rate before the start of the drinking scene, i.e., before the water came out from the tube, indicating that the increased firing rate just before the drinking scene is not due to the olfactory sensory inputs from the water. In summary, while an individual vTT scene cell fires maximally within the scene the cell is in charge of, many of the scene cells begin to increase spiking activity before the actual scene starts.

To further examine the relation of vTT cell firing with scene development, we selected 185 vTT cells that showed clear tuning to one or two specific scenes and aligned these vTT cells by the timing of maximal firing as a function of scene development during the task (Fig. 6a). We found that individual vTT scene cells were tuned maximally to a smaller scale scene or sub-scene within each behavioral scene. For example, different odor checking scene cells were tuned maximally to different sub-scene within the odor checking scene. Distinct drinking scene cells showed maximal tuning to different sub-scene within the drinking scene. Surprisingly, the repertoire of maximal tuning of these vTT cells covered virtually all the continuing series of scenes and sub-scenes of the odor-guided reward-directed behavior (Fig. 6a).

**Figure 6:**
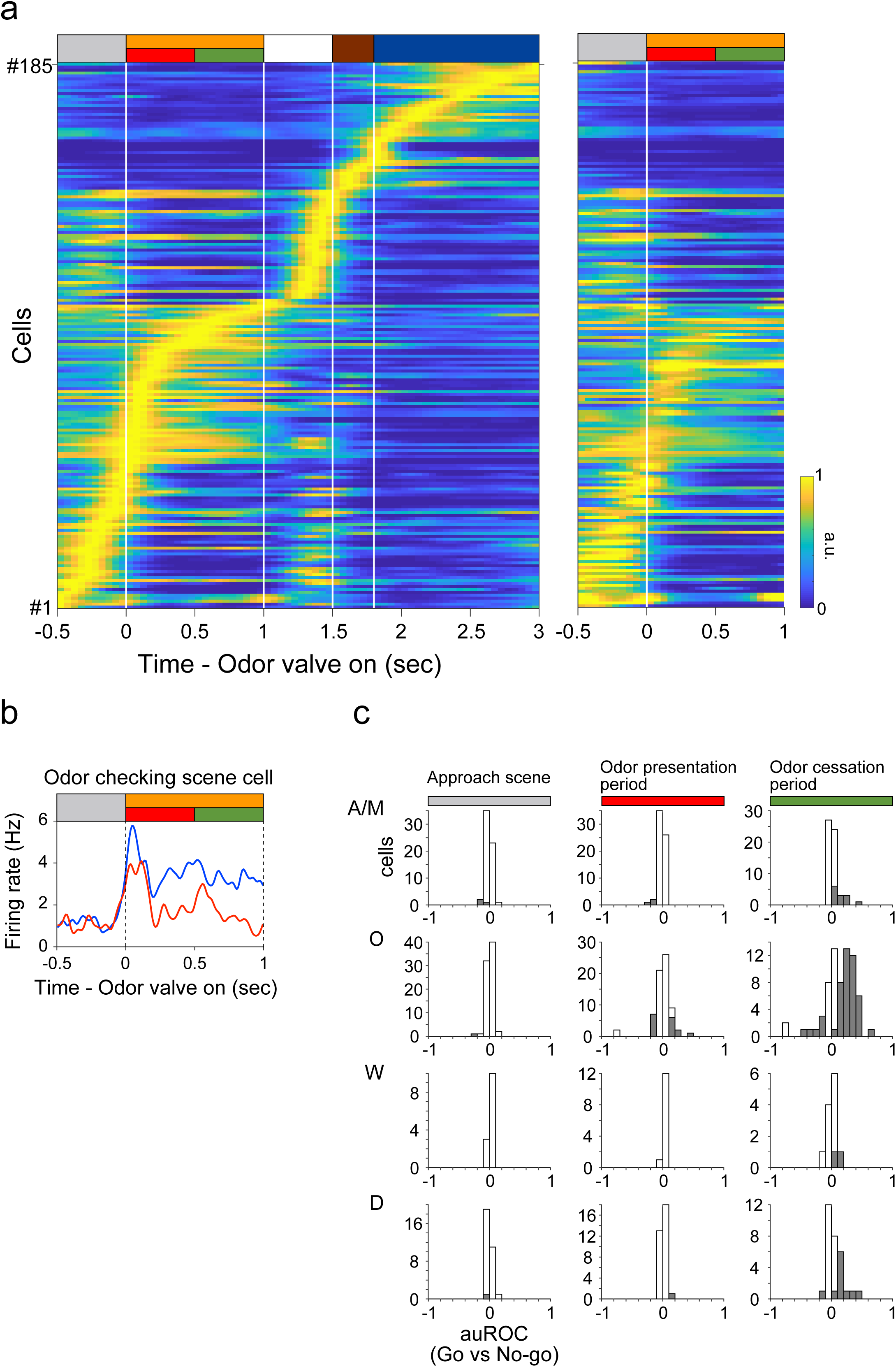
Moment-by-moment functional switching of vTT scene cells during the odor guided go task and the odor guided no-go task. (a) Alignment of vTT scene cells by the timing of maximal firing as a function of scene development during the odor guided go (left) or no-go (right) task. Each bar on the top row shows an individual scene in the task. Gray bar, approaching scene; orange bar, odor checking scene; white bar, moving scene; brown bar, waiting scene; dark-blue bar, drinking scene. The odor checking scene was further subdivided into odor presentation period (red bar) and odor cessation period (green bar). (b) Firing pattern of an odor checking scene cell between go trials (blue line) and no-go trials (red line). (c) Histograms of auROC values of each scene cell during approach scene (gray), odor presentation period (red), and odor cessation period (green) comparing between the go trials and no-go trials. Columns at positive auROC values (> 0) indicate cells that show higher firing rate during go trials, whereas columns at negative values (< 0) indicate cells that show lower firing rate during go trials compared with that during no-go trials. Cells with significant difference in auROC values are filled (t-test, p < 0.05). A/M, approaching and moving scene cells; O, odor checking scene cells; W, waiting scene cells; D, drinking scene cells.

We expected that, when a go-cue odor was presented in the go trials, the odor checking scene consisted of an odor sniffing sub-scene, a proactive sub-scene when the mouse was sniffing the go-cue odor, and a subsequent reward predicting sub-scene, a reactive sub-scene when the mouse was predicting the emergence of reward in the reward port based on the go-cue odor. However, observation of the mouse’s behavior did not allow us to determine the exact timing of the initiation of the reward predicting sub-scene. We also expected that, in the trial in which a no-go-cue odor was presented, the odor checking scene consisted of an odor sniffing sub-scene and a subsequent no-reward predicting sub-scene although we could not determine exact initiation timing of the no-reward predicting sub-scene. Therefore, to examine the relation between the sub-scenes and the firing pattern we divided the odor checking scene into the odor presentation period (red bar in Fig. 6) during which odor sniffing sub-scene may dominate and the odor cessation period (green bar in Fig. 6) during which reward or no-reward predicting sub-scene may dominate.

We compared the firing pattern of the odor checking scene cells between go trials and no-go trials (Fig. 6b, c). During the odor presentation period (red in Fig. 6) only 21.1% of odor checking scene cells showed significantly different firing rate between go trials and no-go trials. In contrast, during the odor cessation period, 61.8% of odor checking scene cells showed significantly different firing rate between them (green in Fig. 6). We thus focused on the odor cessation period. Among the odor checking scene cells that showed significantly different firing rate between go and no-go trials, a majority of cells (87.2%, 41/47 cells) showed higher firing rate during the odor cessation period of go trials compared with that of no-go trials (compare left and right charts in Fig. 6a). Only a small number of odor checking scene cells (12.8%, 6/47 cells) showed higher firing rate during the odor cessation period of no-go trials compared with that of go-trials.

Fig. 6b shows an example of odor checking scene cell showing higher firing rate during go trials compared with no-go trials. This odor checking scene cell started to show higher firing rate in go-trials (blue line) at the odor presentation period and continued the higher firing rate during the subsequent odor cessation period.

These results indicate that the firing pattern of odor checking scene cells differ clearly depending on the odor-guided prediction of the future scene. Many odor checking scene cells showed higher firing rate during the presumptive reward predicting sub-scene after go-odor stimulation whereas they showed lower firing rate during the presumptive no-reward predicting sub-scene after no-go odor stimulation.

We also observed that some approaching and moving scene cells, waiting scene cells and drinking scene cells showed significantly different firing rate between go trials and no-go trials during odor cessation period, whereas they presented no significant change in firing rate during approaching scene and odor presentation period (Fig. 6c). These results suggest that some scene cells start to increase their firing rate at the scene when the mouse is predicting the future scenes that lead to the goal scene of the task (drinking and eating). These results suggest that the activity of scene cells in the vTT represents not only a present scene but also future scenes the mouse predicts (Sharpe & Schoenbaum, 2016).

### Cell types and connection pattern of the vTT

Although a majority of neurons in layer II of the vTT are pyramidal cells, vTT also contains other cell types (Haberly and Price 1978, Nevil and Haberly 2004). To examine the distribution of glutamatergic cells and GABAergic cells in the vTT, we performed *in situ* hybridization for mRNA of vesicular glutamate transporter 1 (VGluT1) and glutamic acid decarboxylase (GAD) 65/67 in the vTT (Fig. 7a, b). About 86% of the vTT cells were *VGlut1*-positive (from three mice), while about 8% of the vTT cells were *GAD65/67*-positive (from three mice). This suggests that principal neurons of the VTT are glutamatergic pyramidal cells.

**Figure 7:**
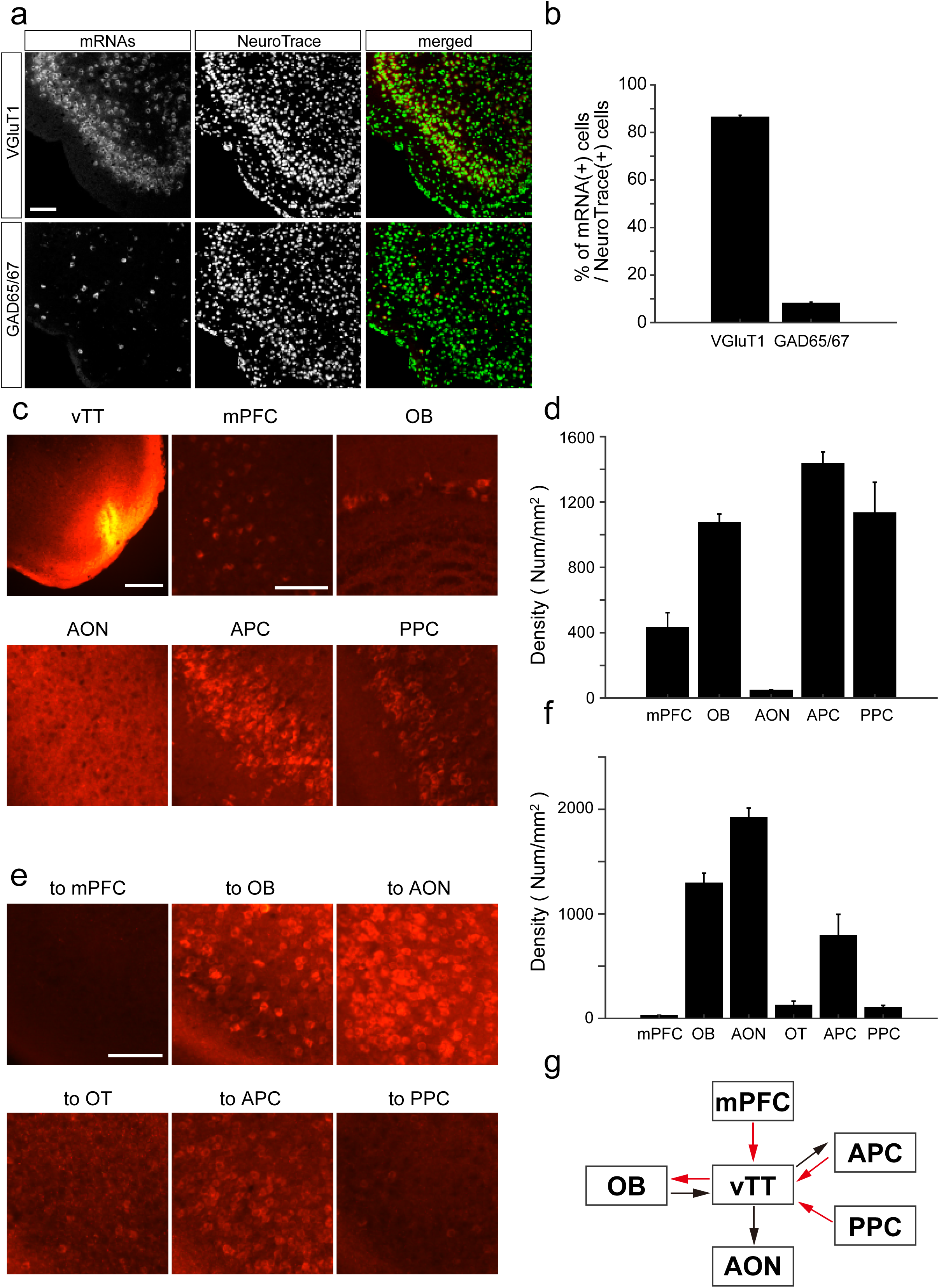
Cell types of vTT neurons and their connections with other areas. (a) *In situ* hybridization for *VGluT1* (upper panels) and *GAD65/67* (lower panels) mRNAs with Neuro Trace staining of vTT cells. Scale bar, 100 μm. (b) Average percentages of *VGluT1* positive cells (left column) and *GAD65/67* positive cells (right column) among the Neuro Trace positive cells in the vTT (n = 3 mice). Error bar, S.E.M. (c) Upper left: Coronal section of the vTT after injection of Alexa 555-conjugated cholera toxin subunit B (CTB, red). Scale bar, 500 μm. The other five panels show CTB-labelled cells after CTB injection in the vTT. mPFC, medial prefrontal cortex; OB, olfactory bulb; AON, anterior olfactory nucleus; APC, anterior piriform cortex; PPC, posterior piriform cortex. Scale bar, 100 μm (d) Average density of CTB-labelled cell bodies in the each area (mPFC, n= 5 from 3 mice; OB, n = 5 from 3 mice; AON, n = 3 from 2 mice; APC, n = 5 from 3 mice; PPC, n = 5 from 3 mice). Error bar, S.E.M. (e) CTB-labelled vTT cells after injection of CTB into the mPFC (upper left), OB (upper middle), AON (upper right), olfactory tubercle (OT, lower left), APC (lower middle) and PPC (lower right). Scale bar, 100 μm. (f) Average density of retrogradely-labelled CTB-positive cells in the vTT (mPFC, n = 3 from 2 mice; OB, n= 3 from 2 mice, AON, n = 3 from 2 mice; OT, n = 5 from 3 mice; APC, n = 4 from 4 mice; PPC, n = 3 from 2 mice). Error bar, S.E.M. (g) Schematic diagram of the connectivity pattern of vTT. Arrows show axonal projection. Black arrows, presumptive afferent connections. Red arrows, presumptive top-down connections.

It has been reported that vTT have reciprocal connections with the olfactory bulb (OB), anterior piriform cortex (APC), and posterior piriform cortex (PPC) (Luskin and Price 1983a,b, Igarashi et al, 2012). In addition, the deep layers of the vTT receive top-down inputs from the medial prefrontal cortex (mPFC) (Hoover and Vertes, 2011). To further examine cortical areas that project axons to the vTT, we injected a retrograde tracer, cholera toxin B subunit (CTB) conjugated with Alexa 555, into the mouse vTT (Fig. 7c). A number of retrogradely-labelled (CTB-positive) cell bodies were identified in the OB, APC, PPC, and mPFC, whereas CTB-positive cell bodies were hardly observed in the anterior olfactory nucleus (AON), which is located just dorsal to the vTT (Fig. 7d). To examine cortical areas that receive axonal projection from vTT cells, we injected CTB into the mPFC, OB, AON, olfactory tubercle (OT), APC, and PPC. We then counted retrogradely labelled CTB-positive cells in the vTT (Fig. 7e, f). Many vTT cells were retrogradely labelled from the OB, AON, and APC, but only a few cells were retrogradely labelled from the OT and PPC. Retrogradely labeled cells were very scarce in the vTT following injection of CTB into the mPFC. These results indicate that, in addition to the heavy reciprocal connection with the OB, the vTT projects axons to the AON and APC and receives top-down projections from the APC, PPC and mPFC (Fig. 7g).

## Discussion

### Scene cells in the vTT

Present results demonstrate characteristic tuning of individual vTT cells of the olfactory cortex to the scene the mouse encounters during the learned feeding and drinking behaviors; individual vTT cells fired maximally whenever the mouse faced to a particular scene of the learned behavior. Because most of recorded vTT cells showed the clear scene-selectivity of their maximal firing, we named these cells “scene cells”.

In the odor-guided eating or no-eating task (Figs 1-3), eating scene cells in the vTT fired maximally during the eating scene, whereas they tended to be silent during the instrumental scene (approaching scene and food deprivation scene). In a striking contrast, instrumental scene cells fired maximally during either the approaching scene or the food deprivation scene. These instrumental scene cells were nearly silent during the eating scene.

In the odor-guided go task to drink water (Figs 4-6), we classified vTT cells into approaching and moving scene cells, odor checking scene cells, waiting scene cells, and drinking scene cells based on their firing pattern during the go trials. Approaching and moving scene cells fired maximally at the scene when either the mouse approached to the odor port or moved from the odor port to the water port. Odor checking scene cells fired maximally at the scene when the mouse examined the odor cue in the odor port. Waiting scene cells fired maximally at the scene when the mouse waited water to come out from the tube, and finally, drinking scene cells fired maximally at the scene when the mouse took the water reward. Therefore, a majority of neurons in the vTT showed maximal firing at a particular scene, suggesting ‘one cell - one scene’ relationship in these cells. However, we also noted that a subset of vTT cells showed highly increased firing rate at two or more different scenes (Fig. 5a, b).

The presence of scene cells suggests critical roles of contextual scene information in olfactory sensory processing in the vTT. vTT cells send the axon to other olfactory cortical areas such as AON and piriform cortex (Fig. 7, Haberly and Price 1978, Luskin and Price 1983). Further experiments are needed to determine whether other areas of the olfactory cortex contain scene cells. It is also of great interest to examine whether the gustatory cortex and the oral area of the somatosensory cortex contain scene cells that fire maximally during a specific scene of the feeding and drinking behaviors.

### Scene-dependent olfactory sensory processing

What are the possible functions of vTT scene cells with respect to olfactory sensory processing? Pyramidal cells are principal neurons in the vTT and extend apical tuft dendrites to the most superficial layer (layer Ia) (Fig. 8) (Haberly and Price 1978). The apical tuft dendrites receive excitatory synaptic input in layer Ia from axon terminals of mitral cells of the olfactory bulb (Friedman and Price 1984, Igarashi et al 2012, Nagayama et al, 2010), and excitatory synaptic inputs in layer Ib from Ib association fibers of other pyramidal cells of the olfactory cortex (Luskin and Price 1983). These synaptic inputs on apical tuft dendrites in layer I may convey olfactory sensory information directly from the olfactory bulb or indirectly after the relay in olfactory cortex areas.

**Figure 8.**
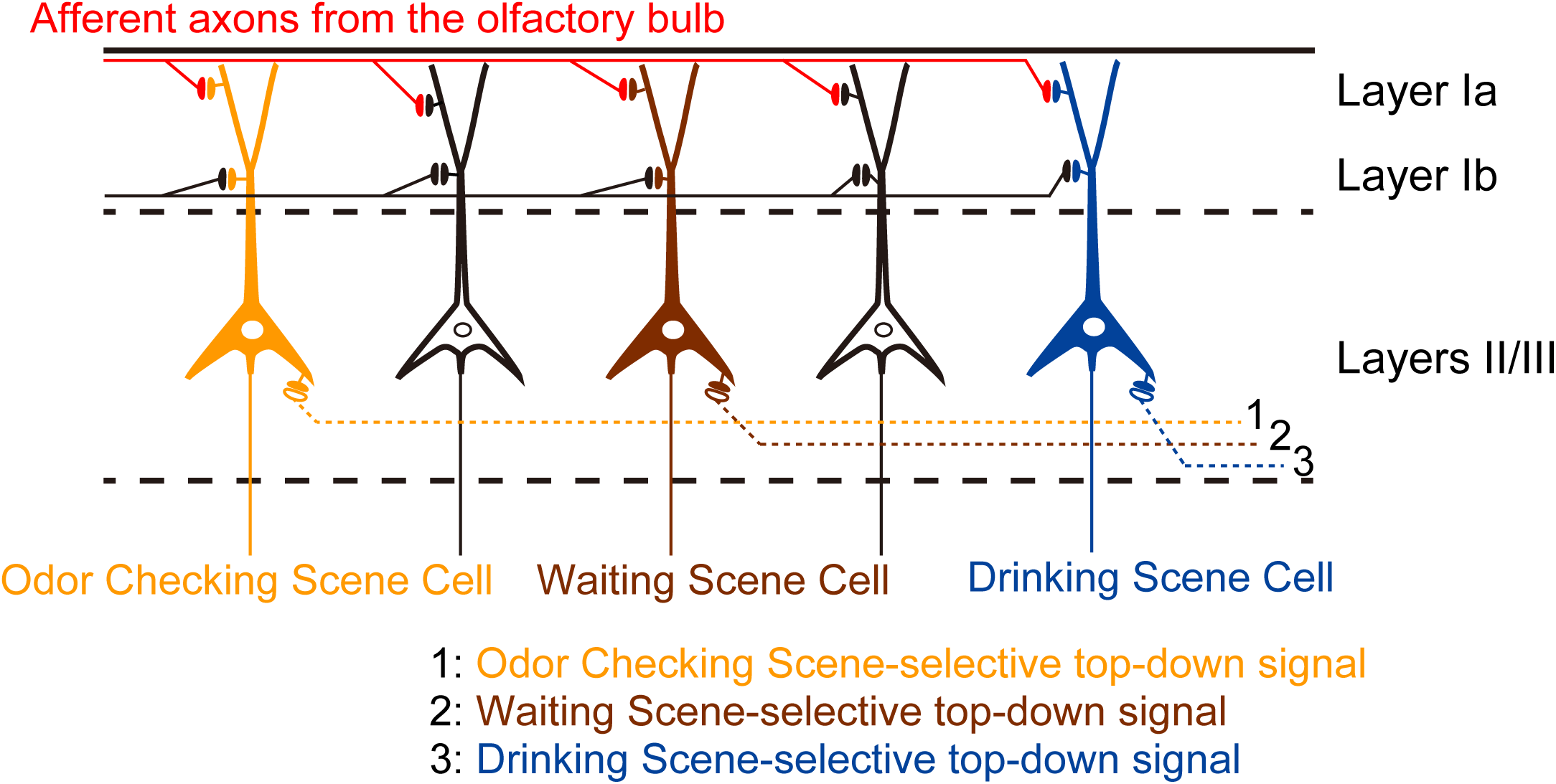
Schematic diagram illustrating the hypothesis that individual vTT pyramidal cells of the olfactory cortex are scene-dependent coincidence detectors, integrating bottom-up olfactory sensory signals with top-down scene-selective signals. For simplicity, this diagram shows only three types of scene cells each with a scene-selective top-down input. Orange shows the odor checking scene cell that receives odor checking scene-selective top-down signal from higher cortical areas such as medial prefrontal cortex. Brown shows the waiting scene cell that receives waiting scene-selective top-down signal. Blue shows the drinking scene cell that receives drinking scene-selective top-down signal. White cells represent other scene cells. Olfactory sensory inputs include olfactory bulb afferent synapses terminating in layer Ia (red) and association fiber synaptic inputs terminating in layer Ib (black) that are originated from other areas of the olfactory cortex.

In addition, vTT pyramidal cells extend basal dendrites and proximal apical oblique dendrites in layers II and III receiving top-down inputs from medial prefrontal cortex (Hoover and Vertes 2011) and top-down deep association fiber inputs from the piriform cortex. Therefore, individual pyramidal cells in the vTT receive olfactory sensory inputs and top-down inputs on spatially well-segregated compartments of dendrites (Fig. 8).

It has been shown that pyramidal cells in the neocortex can detect the occurrence of near-simultaneous synaptic inputs impinging on spatially segregated dendritic compartments and generate action-potential bursts in response to the coincident synaptic inputs (Larkum et al., 1999; Stuart & Spruston, 2015; Hill et al., 2013; Sakmann, 2017). Therefore, neocortical pyramidal cells have the capacity for coincidence detection of spatially separated subthreshold synaptic inputs.

Because pyramidal cells in the vTT have dendritic morphology similar to neocortical pyramidal cells, we speculate that vTT pyramidal cells are capable for coincidence detection of spatially segregated synaptic inputs; i.e., coincident detection of olfactory sensory inputs on apical tuft dendrites in layer I and top-down inputs on basal dendrites and proximal apical oblique dendrites in layers II and III (Fig. 8).

During the approaching scene, approaching and moving scene pyramidal cells may receive top-down excitatory synaptic input in deep layers (layers II and III). If the approaching and moving scene pyramidal cells receive olfactory sensory input in layer I during the approaching scene, the top-down input may augment or multiply responses to the olfactory sensory input. In other words, the approaching and moving scene pyramidal cells may function as coincidence detectors and respond to the two coincident inputs with action-potential bursts during the approaching scene. On the other hand, a majority of odor checking scene cells, waiting scene cells and drinking scene cells do not receive top-down excitatory synaptic input during the approaching scene. Therefore, these scene cells may not respond or show only a weak spike responses to the olfactory sensory input during the approaching scene. Only approaching and moving scene cells can function as coincidence detector during approaching scene.

Similarly, during odor checking scene, only odor checking scene cells can function as the coincidence detector between olfactory sensory input and top-down synaptic input. Only waiting scene cells, and drinking scene cells can function as coincidence detector during the waiting scene and drinking scene, respectively. In this way, different scene cells in the vTT may function as coincidence detector of top-down synaptic input and bottom-up olfactory sensory input only during the corresponding scene. Thus during a particular scene of the learned feeding behavior, olfactory sensory information appears to be handled and processed mainly by corresponding scene cells in the vTT (Fig. 8). In other words, the mode of olfactory sensory processing in the vTT changes moment by moment in scene-dependent manner such that distinct scene cells are selected in each scene and assigned to olfactory sensory processing.

We propose that individual vTT pyramidal cells are scene-dependent coincidence detectors, integrating bottom-up olfactory sensory signals with top-down scene signals only during a particular scene of learned feeding and drinking behaviors (Fig. 8). Top-down scene signals might set vTT in a scene-specific working mode for olfactory sensory processing. As the scenes develop toward the goal (eating or drinking) during the feeding behavior, the top-down signals may instruct moment by moment switching of active vTT cell subpopulations in accord with the current and predicted future scenes. To examine these possibilities in more detail, it is necessary to record afferent synaptic inputs and top-down synaptic inputs using in vivo whole cell patch recordings or to optogenetically manipulate these inputs individually (Land et al., 2014).

### vTT cells also represent future scenes the mouse predicts

In the odor-guided eating or no-eating task, many eating scene cells began to increase their firing rate before the mouse start to eat the food. This observation raised the possibility that the increase in firing rate of eating scene cells during the pre-eating scene was not due to the sensory inputs generated by eating but might be due to the prediction of eating based on the sensory inputs from the cue odor. Furthermore, many instrumental scene cells suddenly increased their firing rate before the food was deprived (Fig. 2b bottom), indicating that the firing rate increase during the pre-deprivation period was not caused by actual food deprivation. We speculate that the mouse noticed the experimenter’s hand coming closer to the food dish and predicted that the food dish will be deprived soon (Fig. 2b bottom) and this prediction of danger caused the vTT instrumental scene cells to rapidly increase the firing rate. Based on these observations we hypothesize that not only the present scene the mouse encounters but also future scene the mouse predicts influence firing activity of vTT cells.

In the odor-guided Go/No-go tasks, many odor checking scene cells showed increased discharges that lasted up to the end of the odor cessation period after the mouse sniffed go cue odor which presumably induced the prediction of water reward. In contrast, after sniffing no-go cue odor which presumably did not induce the reward prediction, these cells showed increased discharges only briefly during the odor presentation period and diminished discharges during the odor cessation period (Fig. 6b). These results corroborate the idea that activity of odor checking scene cells are modulated not only by the signals of present scene that the mouse encounters but also by the future scene that the mouse predicts. We speculate that the reward predicting scene activity of the vTT scene cells is not driven directly by the olfactory sensory afferent input but may be induced by top-down inputs from higher areas because the reward predicting scene activity was induced regardless of the odorants used as go-cue. We speculate that the continued high frequency firing of odor checking scene cells during the odor cessation period reflects top-down scene-predicting signals reflecting the continued attention to the predicted reward scene or working memory of the predicted reward generated in the higher brain regions such as medial prefrontal cortex.

Interestingly, higher firing rate during the odor cessation period after go-odor stimulation compared with the firing rate during the same period after no-go odor stimulation was observed not only in odor checking scene cells, but also in a subset of approaching and moving scene cells, waiting scene cells, and drinking scene cells (Fig. 6c). These results suggest that the presumptive top-down reward-predicting scene signals during the odor checking scene occur not only in odor checking scene cells but also in a small subset of scene cells that are in charge of subsequent series of scenes including the goal scene. We speculate that some scene cells may receive top-down scene-predicting signals when the mouse is predicting the scene before the start of the scene.

Similar pre-scene activity was reported in the hypothalamic Agouti-related-peptide (AgRP) neurons, which are interoceptive neurons receiving energy balance-related hormone signals and are known to drive feeding behaviors. It has been reported that the rapid decrease in AgRP neuron firing occurs at the timing when mice detect food-associated cues, prior to actual ingestion of food (Mandelblat-Cerf et al., 2015).

### vTT scene cells have implications on neuronal mechanisms of top-down attention and odor-scene association memory

Present study does not address the question whether the scene-specific increase in firing rate of vTT cells is induced by top-down inputs from prefrontal cortex and higher areas of olfactory cortex or by bottom-up olfactory sensory inputs. If the scene-selective activity of vTT cells is mediated mainly by top-down signals, the scene cell activity may reflect top-down attention signals. Higher centers might send top-down signals to facilitate activity of attended scene cells in the vTT and thus to facilitate attended scene cells’ response to odor stimuli. It is also possible that higher centers might send top-down signals to suppress activity of un-attended scene cells to suppress un-attended scene cells’ response to distractor odor stimuli. Further works with selective inhibition of top-down input or olfactory sensory input are necessary to examine the functional role of the vTT scene cells in odor-guided feeding and drinking behaviors.

The presumptive scene-specific coincidence detection by vTT pyramidal cells may induce burst spike discharges, resulting in the strengthening and weakening of synaptic connectivity between incoming synapses (both bottom-up and top-down) and vTT pyramidal cells, among vTT pyramidal cells via recurrent axon collaterals, and between vTT cells and their target neurons. If specific scene cells are in charge of odor signal processing during a particular scene, the synaptic plasticity occurs in the circuit of these scene cells. Since a majority of vTT cells are scene cells, the synaptic plasticity of vTT cells responsible for odor memory may occur in scene-specific manner.

Attentional modulation also occurs in human olfactory cortex (Zerano, C et al., 2004). However, it is not known at present whether neurons in human olfactory cortex have mechanisms for scene effect on olfactory sensory processing similar to that found in vTT neurons of the mouse olfactory cortex. Human sensory evaluation studies suggest that odor signals from a same food might be processed differently at different scenes of eating and drinking behaviors. For example, our perception of orthonasal odor from wine during pre-drinking aroma-check scene differs strikingly from the perception of retronasal odor from the same wine during the aroma-burst scene just after the swallowing (Shepherd, 2017). The memory of the orthonasal odor of wine differs also from that of retronasal odor of the same wine. Future experiments might reveal evidence for scene-dependent odor processing and odor memory in the human brain, which may be essential for performing and enjoying eating and drinking behaviors at daily meals.

## Materials and Methods

### Animals

All experiments were performed on male C57BL/6NCrSlc mice (9 weeks old; weighing 20–25 g) purchased from Shimizu Laboratory Supplies Co., LTD., Kyoto, Japan. The mice were individually housed in a temperature-controlled environment with a 13-h light and 11-h dark cycle (lights on at 8:00 and off at 21:00). Food and water were available *ad libitum* until behavioral task started. All experiments were performed in accordance with the guidelines for animal experiments at Doshisha University with the approval of the Animal Research Committee of Doshisha University.

### Behavioral task

For the odor-guided eating or no-eating task (Fig. 1a), mice (n = 6) were placed on a food restriction schedule with daily body weight monitoring to ensure that body mass remained within 80% of prior mass before restriction. In the training session, mice were required to associate odors with sugar (sucrose) rewards. The training was conducted in a plastic cage (38.5 x 33.5 x 18 cm, CLEA Japan Inc., Tokyo, Japan) covered with virgin pulp bedding (SLC, Inc., Shizuoka, Japan) and recording camera in easy soundproof room with a ventilator fan providing air circulation and low level background noise. Mice were presented sugar on a holed Petri dish which contained a filter paper (2×2 cm) soaked with one of odors (40 μl) covered with the bedding at an arbitrary position in the cage. Cue odors were eugenol (TOKYO CHEMICAL INDUSTRY Co., LTD., Tokyo, Japan), vanilla essence (NARIZUKA Corporation, Tokyo, Japan). After the learning of association between dish with odor and sugar reward, mice approached and touched the dish, dug the bedding and showed sugar eating behavior.

One day after the initial training, mice were trained to associate almond odor (almond essence, NARIZUKA Corporation) with aversive consequence (malaise) as follows (Raineki et al., 2009). After the mouse approached the dish with almond odor and ate the sugar on the dish, mice received intraperitoneal injection of 0.5M lithium chloride (LiCl, 0.01 ml/g). After the experience of LiCl injection, mice approached the dish with almond odor but left the dish without eating sugar (no-eating response).

After these trainings, we examined the mice to perform odor-guided eating or no-eating task. We randomly presented sugar on the dish with one of three different cue odors (eugenol, vanilla essence, or almond essence). Almond odor was presented in 20% probability. We presented also powder chow on the dish in trials that were randomly inserted among the above trials. At the timing about 6.0 sec after the mice started to eat, we suddenly deprived the dish. Mice performed a session of behavioral tasks consisting of 40-60 trials in a day.

For the odor-guided Go/No-go task (Fig. 4a), we used a behavioral apparatus that was controlled by Bpod State Machine r0.5 (Sanworks LLC, NY, USA), which are open source control devices designed for behavioral tasks. Our system comprises a custom-designed mouse behavior box (Sanworks) with two nose-poke ports on the front wall in a soundproof box (BrainScience・Idea. Co., Ltd., Osaka, Japan) with a ventilator fan providing air circulation and low level background noise. Each of the two nose-pork ports had white light-emitting diode (LED) and infrared photodiodes. Interruption of the infrared beam generated a Transistor-Transistor-Logic (TTL) pulse signaling the entry of the mouse head into the port. Odor deliver port had a stainless steel tubing connected to a custom-made olfactometer (Uchida and Mainen, 2003). Eugenol (TOKYO CHEMICAL INDUSTRY Co., LTD., Tokyo, Japan) was used as a go cue odor, while and amyl acetate (TOKYO CHEMICAL INDUSTRY Co., LTD.) was used as a no-go cue odor. These odors were diluted to 10% in mineral oil and further diluted 1:10 by airflow. Water reward delivery was based on gravitational flow controlled by a solenoid valve (The Lee Company, CT, USA) connected via tygon tubing to a stainless steel tubing. The reward amount (6 μl) was determined by opening duration of the solenoid valve and regularly calibrated.

For the Go/No-go task, mice (n = 6) were placed on a water restriction schedule with daily body weight monitoring to ensure that body mass remained within 80% of prior mass before restriction. Each trial began by the illumination of LED light at the right odor port that instructed the mice to nose poke into the odor port. A nose poke in the odor port resulted in delivery of one of the two cue odors for 500 msec. Mice were required to sniff the odor and then keep nose poking for 500 msec after the cessation of odor stimulation. Five hundred msec after the cessation of odor stimulation, the LED light was turned off and the mice could withdraw its nose from the odor port. If eugenol odor (go cue odor) was presented, mice were required to move to and nose poke into the left water reward port within 2 sec. At the water port, mice were required to keep nose poking for 300 msec before water delivery began. Then water reward was delivered in 6 μl. If amyl acetate odor (no-go cue odor) was presented, mice were required to restrict entering the water port for 2 sec.

### Electrophysiology

Adult male mice were anesthetized with medetomidine (0.75 mg/kg ip), midazolam (4.0 mg/kg ip) and butorphanol (5.0 mg/kg ip) and implanted with a custom-built microdrive of three or four tetrode in the vTT (2.6 mm anterior to the bregma, 0.4 mm lateral to the midline, 4.0 mm from the brain surface). Individual tetrodes consisted of four twisted polyimide-coated tungsten wires (California Fine Wire, single wire diameter 12.5 μm, gold plated to less than 500 KΩ). Two other screws were threaded into the bone above the cerebellum for reference. These electrodes were connected with an electrode interface board (EIB-18, Neuralynx) on the microdrive. The microdrive array was fixed to the skull with LOCTITE 454 (Henkel Corporation, Düsseldorf, Germany). After completion of surgery, mice received atipamezole (0.75 mg/kg ip) to reverse the effect of medetomidine and permit a reduction of recovery period. Mice also received analgesics (ketprofen, 5mg/kg, ip). Behavioral training resumed at least 1 week after the surgery.

Electrical signals were obtained with either a Cheetah recording system (Neuralynx) or the open-source hardware (Open Ephys). For unit recordings, the signals were sampled at 32 kHz in NeuraLynx and at 30 kHz in Open Ephys and band-pass filtered at 600-6,000 Hz. After each recording, tetrodes were adjusted to obtain new units.

### Data analysis

All data analysis was carried out using built-in and custom-built software in MATLAB 2018a (The Mathworks, Inc., MA, USA).

Task accuracy: In odor-guided eating or no-eating task, accuracy rate was calculate as the average of the percentage that the mouse successfully ate the sucrose in the dish with eugenol or vanilla odor, the percentage that the mouse successfully ate powder chow and the percentage that the mouse didn’t eat the sugar in the dish with almond odor. Mice performed 40-60 trials in each session per day. Each block consisted of 5 trials. In odor-guided Go/No-go task, accuracy rate was calculated as the sum of the percentage of success rate in the go trials and the percentage of success rate in no-go trials in a session. Mice performed up to 600 trials in each session per day. Each block consisted of 20 trials.

Spike sorting: Spikes were sorted into clusters offline on the basis of the waveform energy, peak amplitude and principal component1 from the four tetrode channels by means of an automated spike-separation algorithm KlustaKwik (Kenth Harris). The resulting classification was corrected and refined manually with MClust software (A. D. Redish). Cluster quality was quantified by isolation distance. Clusters with isolation distance under 20 were excluded from analysis.

Spike train analysis: In odor-guided eating or no-eating task, we acquired the timestamps of each event (the start of approaching to the dish, touching the dish, and deprivation of the dish) from frames of recorded movie and they were synchronized with spike data. In the odor-guided Go/No-go task, neural and behavioral data were synchronized by inputting the each event timestamps from the Bpod behavioral control system in the electric signal recordings system. To calculate firing rates during tasks, peri-event time histograms (PETHs) were calculated using 10ms bin width and smoothed by convolving spike trains with a 20-msec wide Gaussian filter (Figs 1d, 2d, 5c, 6b). The firing rate difference between late approaching scene and early eating scene in the odor-guided Go/No-go task was verified by the paired t-test.

To examine the relation of firing rate change of individual vTT cells to the development of behavioral scenes in the behavioral tasks, we calculated PETH during tasks. Because scene durations were different in different trials in the odor-guided eating or no-eating task, we standardized each scene time in each trial to average scene time. We calculated PETH using 50 msec bin width and smoothed by convolving spike trains with a 100 msec wide Gaussian filter. To avoid influence of the firing rate differences, PETHs value were divided by peak firing rate of each cell (maximum value of the PETH). Principal component analysis (PCA) was calculated by the singular value decomposition of the normalized PSTHs. Hierarchical clustering was done using the first three PCs of the normalized PSTHs using a Euclidean distance metric and average agglomeration method.

To determine whether a cluster showed a significant scene-specific activity, we used one-way ANOVA with Tukey’s *post hoc* test on a cell-by-cell firing rates in the cluster during the each scene. The area under the receiver operating characteristic (auROC) curves was calculated by comparing the distribution firing rate of each scene across go trials in 50 msec bins to the distribution firing rate across no-go trials. All data are presented as mean ± SEM.

### Histology

After recording, mice were deeply anesthetized by intraperitoneal injection of sodium pentobarbital. Electric lesions were made using 10-20 μA direct current for 5 s to through one of the four leads of tetrode. Mice were perfused transcardially with phosphate-buffered saline (PBS) followed by 4% paraformaldehyde (PFA). Brains were removed from the skull and post-fixed in PFA. The brains were cut in 50 μm coronal sections, and stained with cresyl violet. The position of electrode tracks was determined in reference to the atlas of Paxinos and Watson (2001).

### In situ hybridization

DIG labeled RNA probes for VGluT1 and GAD65/67 were made using the in vitro transcription kit (Roche) according to the manufacturer’s protocol with plasmids kindly provided by Drs. Katsuhiko Ono and Yuchio Yanagawa (Asada, H. et al.,1997; Makinae, K. et al.,2000; Ono, K. et al., 2008). Brain sections were made with a thickness of 20 μm, mounted on slide glasses (Matsunami, CREST) using a paint brash, dried overnight in a vacuum desiccator. The dried sections were fixed in 4% PFA, digested with Proteinase K (10 μg/mL) for 30 min, and post-fixed in 4% PFA. After prehybridization, the sections were incubated overnight at 65°C with DIG-labeled RNA probes. After stringent washing, the sections were blocked with 1% blocking reagent (11096176001, Roche) in TNT for 1 h. Subsequently, the sections were incubated overnight at 4°C with alkaline phosphatase-conjugated anti-DIG antibody (1:1,000; Roche). The sections were washed three times in TNT and once in TS 8.0 (0.1 M Tris-HCl, pH 8.0, 0.1 M NaCl, 50 mM MgCl2), and then alkaline phosphatase activity was detected using an HNPP fluorescence detection set (11758888001, Roche) according to the manufacturer’s instructions. Incubation for this substrate was carried out for 30 min, repeated a total of 3 times, and stopped by washing in PBS. The Sections were counterstained with NeuroTrace Green (Thermo Fisher Scientific) and mounted in PermaFluor (Thermo Fisher Scientific).

### Retrograde tracing

Adult male mice were anesthetized with medetomidine (0.75 mg/kg ip), midazolam (4.0 mg/kg ip) and butorphanol (5.0 mg/kg ip), and then placed in a stereotaxic apparatus (Narishige, SR-5M). Injections were conducted with a syringe pomp (WPI, UltraMicroPump III) connected to a Hamilton syringe (Hamilton, RN-1701) and mounted glass micropipette with a tip diameter of 50 μm connected by an adaptor (Hamilton, 55750-01).

We unilaterally or bilaterally injected 300 nl of CTB conjugated Alexa 555 (Thermo Fisher) at 100 nl/min in mPFC (A/P, 2.4 mm; M/L 0.4 mm from bregma; D/V, 1.0 mm from brain surface), OB (A/P, 4.3 mm; M/L 0.8 mm from bregma; D/V, 1.5 mm from brain surface), AON (A/P, 2.8 mm; M/L 1.3 mm from bregma; D/V, 2.6 mm from brain surface), OT (A/P, 1.5 mm; M/L 1.0 mm from bregma; D/V, 4.7 mm from brain surface), APC (A/P, 2.3 mm; M/L 1.8 mm from bregma; D/V, 3.4 mm from brain surface), and PPC (A/P, −1.5 mm; M/L 3.6 mm from bregma; D/V, 4.5 mm from brain surface) or 250 nl in vTT (A/P, 0.3 mm tilted 30°; M/L 0.4 mm from bregma; D/V, 4.6 mm from brain surface). After the surgery, mice received atipamezole (0.75 mg/kg ip) and ketprofen (5mg/kg, ip). One week later, the mice were deeply anesthetized and perfused with saline and then 4% paraformaldehyde under anesthesia. Brains were cut in 50 μm coronal sections.

### Microscopy

Sections were examined with a confocal laser microscope (Olympus, FV1200), and a bright-field and fluorescent microscope (Zeiss).

## Acknowledgments

We thank the members of the Sakurai lab for valuable discussion. And we thank Hideki Tanisumi for providing illustrations in Figs 1a and 4a. K.S. was supported by JSPS KAKENHI Grant Numbers 18J21358, H.M. was supported by the Takeda Science Foundation, Narishige Neuroscience Research Foundation, and JSPS KAKENHI Grant Numbers 25135708, 16K14557. Y.S. was supported by JSPS KAKENHI Grant Numbers 16H02061.

